# Recombinant Human ACE2-Fc : A promising therapy for SARS-CoV2 infection

**DOI:** 10.1101/2022.07.30.501940

**Authors:** P.K Smitha, R.K. Shandil, Pushkarni Suresh, Kunal Biswas, G.R. Rudramurthy, C.N. Naveenkumar, K. Bharathkumar, Naga Puspha Battula, Suprabuddha Datta Chowdhury, Sakshi Sinha, Sarmistha Dutta, Sujan K. Dhar, Shridhar Narayanan, Manjula Das

## Abstract

SARS-CoV2 entry is mediated by binding of viral spike-protein(S) to the transmembrane Angiotensin-Converting Enzyme-2 (ACE2) of the host cell. Thus, to prevent transmission of disease, strategies to abrogate the interaction are important. However, ACE2 cannot be blocked since its normal function is to convert the Angiotensin II peptide to Angiotensin(1-7) to reduce hypertension. This work reports a recombinant cell line secreting soluble ACE2-ectopic domain (MFcS2), modified to increase binding and production efficacy and fused to human immunoglobulin-Fc. While maintaining its enzymatic activity, the molecule trapped and neutralized SARS CoV2 virus *in vitro* with an IC_50_ of 64 nM. *In vivo*, with no pathology in the vital organs, it inhibited the viral load in lungs in SARS-CoV2 infected Golden-Syrian-hamster. The Intravenous pharmacokinetic profiling of MFcS2 in hamster at a dose of 5 mg/Kg presented a maximum serum concentration of 23.45 µg/mL with a half-life of 29.5 hrs. These results suggest that MFcS2 could be used as an effective decoy based therapeutic strategy to treat COVID19. This work also reports usage of a novel oral-cancer cell line as in vitro model of SARS-Cov2 infection, validated by over expressing viral-defence pathways upon RNA-seq analysis and over-expression of ACE2 and TMPRSS upon growth in hyperglycaemic condition.

## 1. Introduction

Since December 2019, millions of deaths have been reported globally due to the COVID-19 pandemic triggered by Severe Acute Respiratory Syndrome Coronavirus 2 (SARS-CoV2) infection which continues to cause social disruption and pose a significant threat to public health [1, 2]. Several therapeutic strategies and preventive measures such as developing vaccines, neutralizing monoclonal-antibody combinations such as Bamlanivimab/Etesevimab [3], convalescent plasma therapy[4–6], oligonucleotides against SARS-CoV2 genome [7], and repurposing of currently available drugs [8–10] were applied to combat COVID19. Remdesivir [11] was the first repurposed drug used to treat COVID-19 patients while others like Molnupiravir [12], Paxlovid [13] and baricitinib have found entry to the clinic subsequently [14]. However the emergence of multiple variants of the virus from B.1.1.7 (Alpha strain), B.1.351 (Beta strain), B.1.617.2 (Delta strain) to P.1 and B.1.1.529 (Gamma and Omicron strains) [15], calls for the development of alternative therapeutic strategies. Among the proposals for new Covid19 therapies, approaches to neutralize the virus has been explored amongst which drugs targeting the SARS-CoV2 entry receptor, ACE2 (angiotensin converting enzyme II receptor) has gained paramount importance [4,15–20].

The mode of entry of SARS-CoV2 into the host system is through docking of it’s spike receptor binding domain (RBD) to its functional receptor ACE2, present in host cell membrane[15, 21]. ACE2 is a membrane-bound carboxypeptidase known for its activity against vasoactive peptides including angiotensin II (AngII) and Apelin-13, among others [16]. ACE2 receptor exerts its regular function by catalysing the conversion of Ang II to Angiotensin (1-7) which counterbalances the actions of ACE and Ang II, facilitating vasodilation in renin-angiotensin system (RAS) [22]. The enhanced infectivity and severity of SARS-CoV2 infection as compared to earlier coronavirus editions has been attributed to modifications in the viral genome leading to increased affinity towards ACE2 receptor [15, 23]. Thus, blocking of RBD-ACE2 interactions by chemicals, drugs, antibodies and recombinant ACE2 receptor would abrogate binding of virus to host cells and prevent infection. Such an approach has tremendous potential for preventing infection and spread of virus from person to person at early stages [17, 18].

While ACE2 receptor facilitates entry of the virus into the host system, a soluble human recombinant ACE2 receptor molecule is known to inhibit SARS-CoV2 infection by presenting as a decoy receptor [19]. Potency of soluble ACE2 receptor as a therapeutic against SARS-CoV2 infection has been demonstrated in human organoid culture [24] and also in clinical trials [25]. Usage of exogenous ACE2 receptor analogue to neutralize the viral infection is not vulnerable to mutations in the virus as long as it can bind to the ACE2 receptor. In case the virus mutates to decrease the binding affinity to the ACE2 receptor, then it would ultimately result in diminished infectivity[17, 19]. However, soluble ACE2 has been reported to have a shorter circulating half-life, and multiple studies have reported modification of this receptor by expressing it as a fusion protein with human IgG-Fc [15– 18,20,26], either by retaining or abrogating the catalytic activity of the ACE2 protein.

Neutralisation of the virus by affinity-optimized ACE2 receptor variants has been achieved by introducing mutations in ACE2 receptor domains interfacing the spike protein so as to increase the binding strength compared to ACE2 receptor present on host membrane [17,27–29]. Several in vitro studies were conducted to determine efficacy of affinity-optimized variants of ACE2 and ACE2-Fc to potentially block SARS-CoV2 infection[27, 29]. Most of these molecules were tested in 2D cellular models and their in vivo activities are yet to be fully established. Nevertheless, these studies have established the proof of concept of prophylactic or therapeutic efficacy of ACE2 molecule.

Here, we report the design and development of recombinant human ACE2 receptor conjugated to Fc domain of hIgG molecule (Ace2Fc) and a mutated version of the same (MFcS2) both of which serve as decoy receptors for SARS-CoV2. Further *in vitro* comparison demonstrates binding affinity of MFcS2 to be superior than Ace2Fc. We report results of extensive characterization of MFcS2 including its Circular Dichroism spectra, in vitro peptidase activity, in vitro and in vivo efficacy and pharmacokinetics properties. Further, in addition to Vero-E6, which is an African green monkey kidney epithelial cells widely used to demonstrate the in vitro neutralization of anti-SARS CoV2 therapies [30–32], we have established a human oral cell line model for SARS-CoV2 infection, since oral cavity is among the main route of entry for this virus into the human system. Results presented in this manuscript demonstrates neutralization of SARS-CoV2 by MFcS2 in the oral cell line model indicating potential of the molecule to be developed as formulations to block entry of the virus through oral and nasal cavities. The MFcS2 protein will not only help lower the infection rate but being enzymatically active, will also be available for the breakdown of Ang II peptide, thereby keeping a check on proper regulation of the renin-angiotensin pathway. This molecule exhibited good efficacy, encouraging pharmacokinetics and therapeutic potential in a Golden Syrian hamster model of SARS-CoV2 infection. These results suggest that MFcS2 can be developed as promising therapeutic candidate for treatment of COVID-19.

## 2. Materials and Methods

### 2.1 Materials

ACE2 cDNA ORF clone (HG10108-ACR) was purchased from Sino biological, China. SARS-CoV2 strain (USA-WA1/2020) used in this study was sourced from BEI Resources, USA. SARS-COV2 spike-1 protein (P-104) and Goat anti-human IgG HRP conjugate (C-002) were procured from Cognate Biolabs, India. Other critical reagents used in the study included Lipofectamine™ 3000 Transfection Reagent (L3000-001), Penicillin-Streptomycin (15140-122), Fetal Bovine Serum (10270106), CyQUANT™ LDH Cytotoxicity Assay kit (C20301) and Zeocin (46-0509) from ThermoFisher scientific, Zinc chloride(Z0152) and DMEM (D5648) from Sigma-Aldrich, RPMI-1640 (AT028A) from HiMedia, Mca-APK(Dnp) (BML-P163) from Enzo life science and RT-PCR Detection Kit (CM4R01) and RNA Extraction Kit (NDX-RNA-021) from NeoDx.. All the other reagents used were commercial reagent-grade products.

### 2.2 Animals

Golden Syrian hamsters were procured from certified animal vendors and housed in individually ventilated cages (IVCs). Feed and water were provided *ad libitum*. Animal husbandry and experimental protocols were reviewed and approved by the Institutional Animal Ethics Committee (IAEC) of FNDR (IAEC,2082/PO/RC/S/19/CPCSEA), registered with the Committee for the Purpose of Control and Supervision (CPCSEA), Government of India.

### 2.3 Cell culture

HEK-293E and Vero-E6 cell lines were cultured in DMEM and MhCT08-E [33] oral squamous cell carcinoma epithelial cell line was cultured in RPMI-1640 containing 11.1 mM and 25 mM glucose, as appropriate. All the media used were supplemented with 10% FBS and 1X Penicillin-Streptomycin and cells were cultured at 37 °C in 5% CO_2_ in air.

### 2.4 Constructs

IgGFc: Human IgG Fc Sequence (Supplementary Material Figure S1) was codon optimized for HEK293 expression and synthesized from Genscript USA in pUC57 vector. pTT5-hFc clone was constructed by amplifying the IgG-Fc region from pUC57-IgG-Fc with primers IgG-Fc forward and reverse (Supplementary Material Table S1) and ligating into NheI-BamHI sites of pTT5 vector.

Ace2Fc: pTT5-ACE2Fc was constructed by amplifying the ectodomain sequence corresponding to a.a 1-740 (NM_021804.1) from ACE2 cDNA ORF clone (HG10108-ACR, Sino biological) with primers Ace2 forward and reverse (Supplementary Material Table S1) and inserting it into PmeI-NheI sites of pTT5-hFc clone.

MFcS2: Modified human ACE2 (Sequence-Supplementary Material S1) was codon optimized for HEK293 expression and synthesized from Genscript USA in pUC57 vector. Further it was amplified with primers MFcS2 forward and reverse (Supplementary Material Table S1) and cloned into EcoR1-NheI sites of pTT5-hFc to make pTT5-MFcS2 clone. MFcS2 sequence was subcloned into a modified pCDNA3.1 Zeocin vector (Data not shown) between EcoRI-BamH1 site to make pCDNA3.1-MFcS2.

### 2.5 Transfection

For transient transfection, pTT5-ACE2Fc and MFcS2 clones were transiently transfected into HEK293E cells using Lipofectamine 3000 reagent as per manufacturer’s instructions. Briefly, 3X10^6^ HEK293 E cells were transfected with 24 µg of plasmids and 22 µL of Lipofectamine and 48 µL of P3000 reagent diluted in Opti-MEM medium and incubated for 48 to 96 hours.

### 2.6 Generation of MFcS2 expressing stable cell line

For generation of HEK293E cells stably overexpressing the MFcS2 protein, the pCDNA3.1-MFcS2 construct was transfected using Lipofectamine 3000 as per manufacturer’s instruction. Cells were grown in 400 µg/mL zeocin for three weeks and surviving cells were counted, diluted and re-plated in a fashion that there is one cell per well of a 96 well plate. Expression of secreted MFcS2 protein of each clone was confirmed by analysis of the conditioned medium in ELISA. The highest expressing clone was chosen for further growth and protein production.

### 2.7 Protein Purification

The conditioned supernatant containing ACE2-Fc or MFcS2 protein was bound to protein-A affinity beads. The column was washed with 20mM sodium phosphate buffer pH 7.4 and eluted with 0.1M glycine buffer containing 150mM NaCl, pH 2.7. The eluted protein was instantly neutralized with 1M Tris-HCl, pH 8.0 and stored in formulation buffer containing 100 mM glycine, 150 mM NaCl, and 50 μM ZnCl2 at a pH of 7.5.

### 2.8 Electrophoresis and Western blotting

The purified proteins were analysed on 8% sodium dodecyl sulfate-polyacrylamide gel electrophoresis (SDS-PAGE) under non-reducing and reducing conditions. Protein samples were transferred onto a Nitrocellulose membrane and blocked with 5% skim milk in 1xPBST (0.05% Tween20) and probed with anti-human IgG-HRP for 1 hour at 37°C.

### 2.9 Enzyme Linked Immunosorbent Assay (ELISA)

To compare the binding affinity of ACE2-Fc and MFcS2 protein, immunosorbent plate was coated with 1 µg/mL of Spike protein overnight at 4°C. Varying concentrations of ACE2-Fc and MFcS2 (2.5µg/mL to 0.625 µg/mL) were incubated for 1 hour at 37°C followed by the addition of anti-human IgG-HRP. Reaction was developed with TMB peroxidase substrate and the absorbance was recorded at 450 nm on a microplate reader.

### 2.10 Secondary structure determination by Far-UV CD spectroscopy

Far-UV circular dichroism (CD) spectra data of purified MFcS2 protein solution in 20mM phosphate buffer pH 7.4 at 25°C, were collected from a Jasco J-815 spectrometer using a 2-mm pathlength quartz cuvette over wavelength range of 200 to 250 nm. Spectra were analysed using the K2D tool [34] for abundance of secondary structure elements.

### 2.11 Cytotoxicity assay

Levels of lactate dehydrogenase (LDH) enzyme released in cytosol is a standard marker of cellular cytotoxicity. Vero-E6 monolayers were incubated with MFcS2 for 48 hours at 37^°^C in DMEM following which cytosolic LDH level was determined using CyQUANT™ LDH Cytotoxicity assay kit following kit manufacturer’s protocol.

### 2.12 In vitro neutralization assay by Plaque Reduction Neutralization Test (PRNT)

SARS-CoV2 virus was propagated in Vero-E6 cells. For PRNT assay, six serial dilutions of MFcS2 (500 µg/mL to 15.625 µg/mL) were mixed with ∼30 PFU/well of virus and were added to Vero-E6 and MhCT08-E cells followed by incubation for 1 h at 37 °C and 5% CO_2_ in a 96 well plate. Inoculum was removed subsequently and replenished with DMEM. Cells were harvested 48 hours post infection for qRT-PCR, or were overlayed with equal volumes of DMEM with 4%FBS and 2% carboxy methylcellulose (CMC) for plaque reduction neutralization test (PRNT). Plates were further incubated for 72 hours at 37 °C in a CO2 (5%) incubator. Cells were then fixed with paraformaldehyde and stained with crystal violet to count the plaques visually. Number of plaques reduced in the MFcS2 treated with respect to virus infected cells were determined, and percentage reduction was calculated in each tested concentration of MFcS2. IC_50_ of MFcS2 against SARS-CoV2 was determined by fitting the dose-concentration values to a 4-parameter logistic regression equation using Graphpad Prism version 9.0.

### 2.13 Real time PCR assay

Total RNA was extracted using RNA extraction kit as per manufacturer’s instruction. Viral entry in MFcS2-treated and untreated Vero-E6 cells or hamster lungs were determined by quantitative real-time reverse transcription PCR (qRT PCR) assay using SARS-CoV2 RT-PCR detection Kit as per manufacturer’s instructions. Gene copy number of N (Nucleocapsid), E (Envelope) and RdRp (RNA dependent RNA polymerase) gene of SARS-CoV2 were determined in RNA extracted from cell and tissue lysate, as appropriate. ACE2 and TMPRSS expression levels in MhCT08-E cells grown in low and high glucose media were checked with gene-specific primers (Supplementary Table S2).

### 2.14 RNA sequencing of virus infected cells

Transcriptome sequencing for the extracted RNA from SARS-CoV-2 infected MhCT08 cells was performed at Eurofins Genomics, Bangalore after checking the quality of RNA on Qubit Fluorometer followed by Agilent 4200 Tape Station. Libraries were prepared from the QC-passed RNA samples using illumina TruSeq Stranded mRNA sample Prep kit and PCR enriched libraries were further checked on Agilent 4200 Tape Station. Libraries were sequenced on illumina NextSeq500 platform using 150 bp paired-end chemistry. Reads obtained from the sequencer were quality checked and filtered using fastp [35] and subsequently aligned using STAR [36] pipeline. Differential gene expression analysis for the extracted counts were carried out using DESeq2 R package [37] and over-representation of pathways in differentially expressed genes were analyzed using clusterProfiler[38].

### 2.15 Pharmacokinetics

For intravenous (IV) pharmacokinetic studies, hamsters were injected intravenously via jugular vein with 5 mg/kg MFcS2 at a concentration of 2mg/mL. Blood samples were collected at 1min, 15 min, 2h, 8h, 24h and 48h post administration from two animals per time point. For intranasal study, hamsters were anesthetized with Ketamine Xylamine IP injection to prevent reflex and were gently placed on a tray in supine position in biosafety cabinet. With a micropipette 50 µL of 2.5 mg/mL MFcS2 (2.5mg/kg body weight) was delivered per nostril. Animals were allowed to recover on the tray for 30 minutes to allow diffusion of the protein into lungs. Two hamsters were euthanised at each time point (1h, 8h, 24h and 48 hours post dosing) and terminal blood and lung tissues were harvested. MFcS2 concentrations were determined by a custom ELISA developed in-house in serum separated from blood samples and in lung tissue homogenate prepared by homogenizing the tissue in a volume of 1X PBS equivalent to the weight of the tissue and clarifying by centrifugation and collection of supernatant. For ELISA, anti-Human IgG was used as capture and corresponding HRP conjugate was used as detection with ACE2-Fc as standards.

Pharmacokinetics parameters including area under the curve (AUC), maximum plasma concentration (C_max_), time to reach maximum plasma concentration (T_max_), terminal half-life (t_1/2_), and clearance were evaluated by a non-compartmental analysis method using PKsolver2, (version 2.0) plugin in Microsoft Excel [39].

### 2.16 Safety assessment

Six vital organs (Lungs, liver, kidney, spleen and heart) from hamsters used in pharmacokinetic study were harvested for histopathological examination. Organs were fixed in 10% neutral buffered formalin, mounted on slides and stained with hematoxylin/eosin. Gross pathology like oedema, inflammation, haemorrhages, and histopathological features such as, interstitial pneumonia, diffuse alveolar damage, cellular infiltration, necrotic cellular debris and mononuclear cells were scored.

### 2.17 In vivo neutralization assay

To check the *in vivo* neutralization potential of MFcS2, hamsters were deeply anaesthetised as described in section 2.15 and infected with 5×10^5^ plaque forming units (PFU) per lung of SARS-CoV2 intranasally. Briefly, a 100 µL of virus preparation containing 5×10^5^ PFU was taken into micropipette and 50 µL was dropped into each nostril of the hamster followed by recovery of 30 minutes in upright position. Infected animals were randomized into control and treatment groups. Hamsters in the treatment group were dosed with MFcS2 (5mg/kg = 250µg/100 µl/hamster) intravenously via juglar vein. One animal each from both the groups were euthanized at 0,1,2,6,12,24,48 and 96 hrs by CO_2_ narcosis and the lungs were examined for any gross pathology. One half of the lung tissue was processed for qRT-PCR as described in section 2.13 and other half was homogenised with equivalent volume of phosphate buffered saline. The homogenate was clarified by centrifugation and used for enumerating the viral particles by PFU as described in section 2.12. PFU counts were normalized for per gram of lung tissue.

### 2.18 Measurement of Peptidase activity of MFcS2

Hydrolysis of (7-methoxycoumarin-4-yl) acetyl-Ala-Pro-Lys(2,4-dinitrophenyl) (Mca-APK-DNP; Enzo Life Sciences) was used to quantify MFcS2 peptidase activity. Two dilutions of MFcS2 (2.5 and 5.0 nM) was incubated in 100 mM Tris-HCl, 1 M NaCl, 600 μM ZnCl_2_(pH 7.5) containing 50 μM of Mca-APK(Dnp) at 37^°^C for 20 minutes in a reaction volume of 100 μL. An increase in fluorescence (320 nm excitation/430 nm emission) was monitored over time using Synergy^TM^ H1multi-mode reader (BioTek).

## 3. Results

### 3.1 MFcS2 protein showed improved binding affinity to SARS-CoV2 Spike protein

Both proteins were produced in HEK293E cells either by transient transfection for ACE2-Fc or from the stably expressing MFcS2 clone (Figure 1a), purified using protein A beads and were identified on a reduced SDS-PAGE gel with molecular weights at ∼130 kilodalton (kDa), which was consistent with the predicted molecular weight of their monomers (Figure 1b). Western blot analysis using IgG specific antibody further showed a single band at ∼130 kDa under reducing condition and ∼250 kDa under non-reducing conditions (Figure 1c and 1d). The binding specificity of ACE2-Fc and MFcS2 to SARS-CoV2 spike protein was confirmed by enzyme linked immunosorbent assay (ELISA). MFcS2 showed a higher binding affinity to the spike protein of SARS-CoV2 when compared to ACE2-Fc protein (Figure 2), hence was taken forward for further studies.

**Figure 1:**
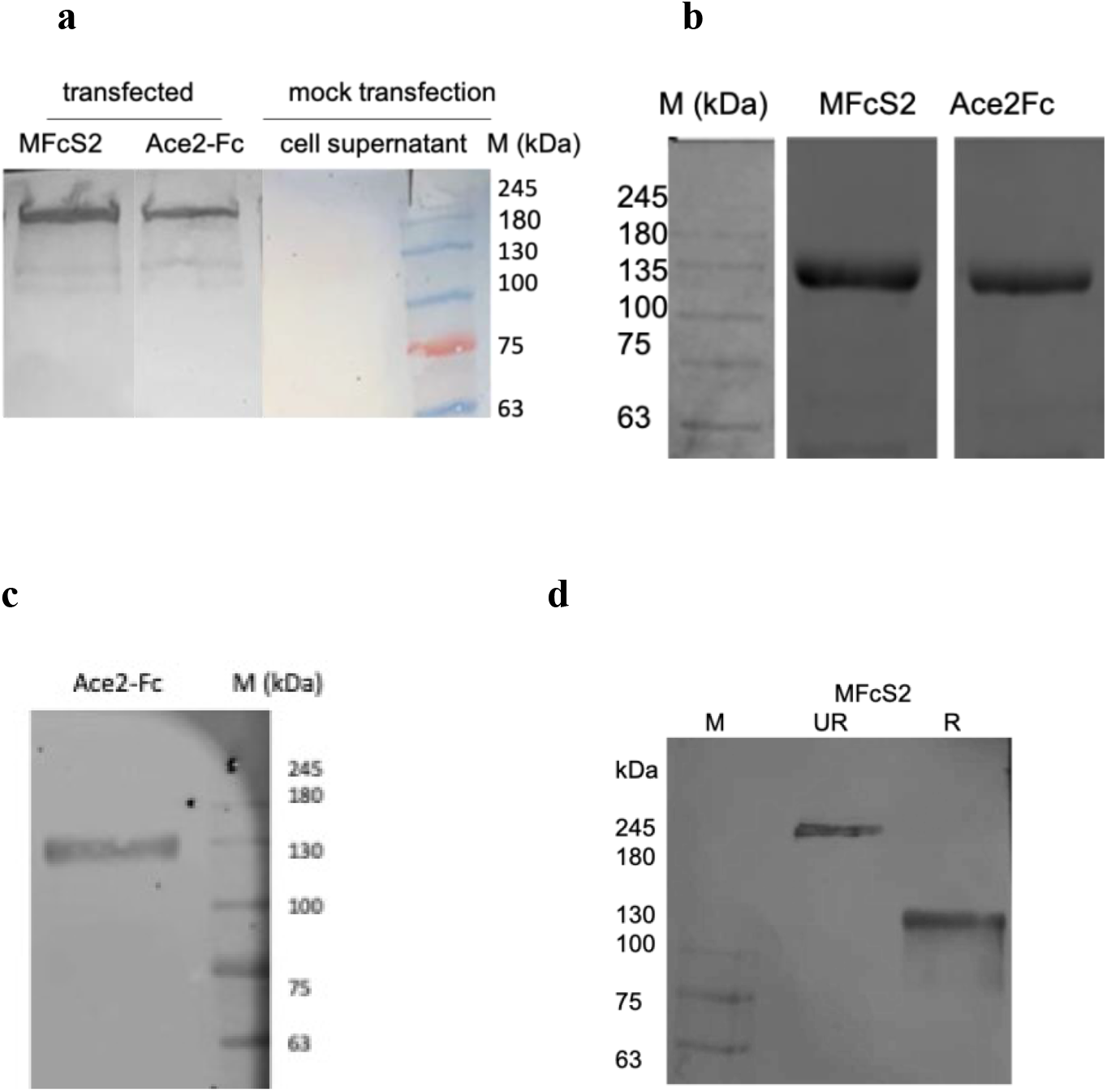
PAGE and Western analysis of Ace2-Fc and MFcS2. **a:** Western blot analysis of MFcS2 and Ace2-Fc from conditioned supernatants from transient transfections with and without the plasmids in non-reducing condition **b:** SDS-PAGE analysis of purified MFcS2 and Ace2-Fc in reducing condition **c:** Western blot of purified Ace2-Fc in reduced condition **d:** Western blot of unreduced (UR) and reduced (R) MFcS2 protein purified from stably expressing HEK293E_MFcS2 cell line. The blots were probed with Goat anti-Human HRP and developed with DAB/H2O2

**Figure 2:**
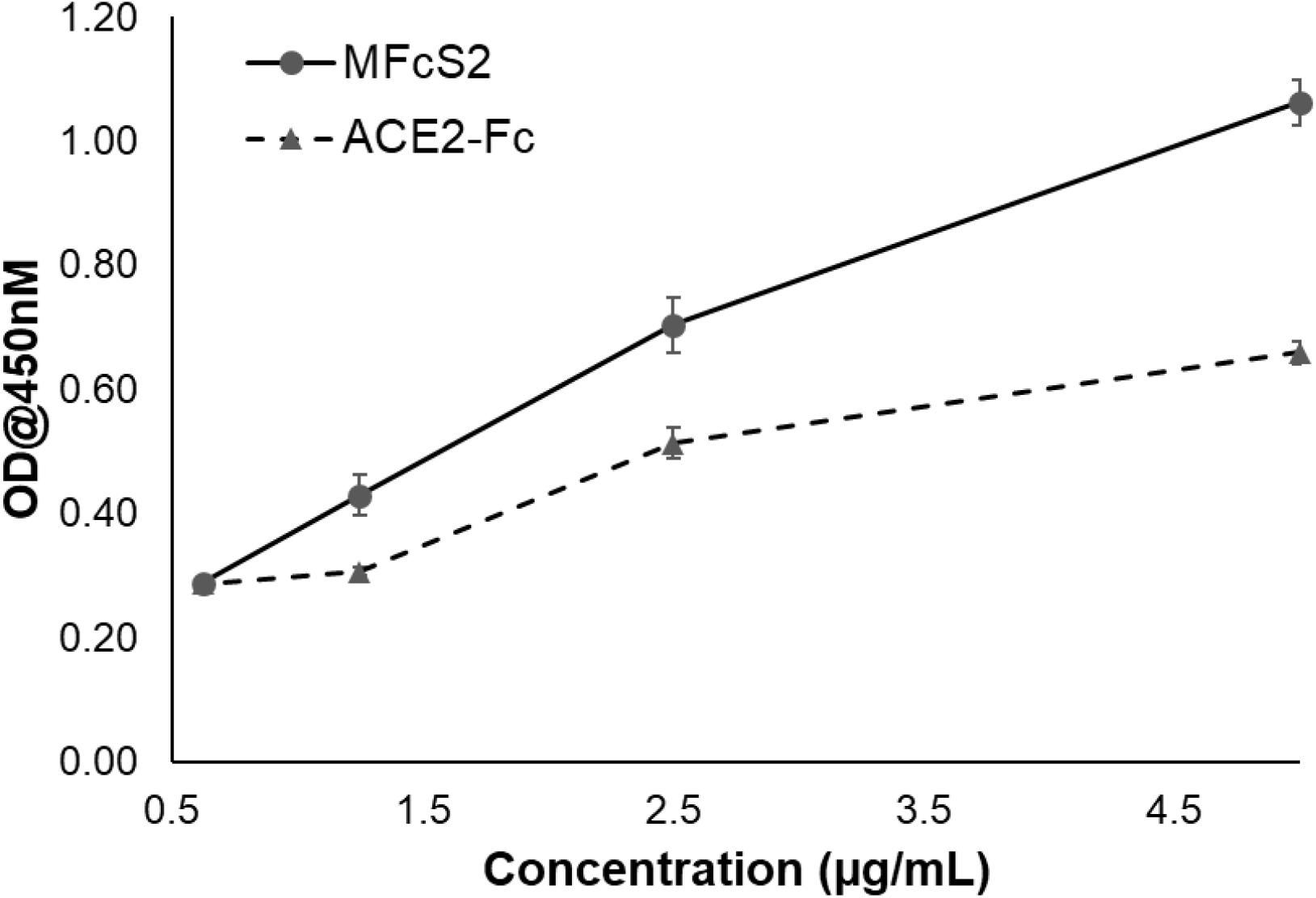
Binding affinity of MFcS2 and Ace2-Fc protein to the SARS-CoV2 Spike protein demonstrated by ELISA

### 3.2 Secondary structure analysis of MFcS2

Far UV CD spectral analysis showed that MFcS2 consists of 46% alpha helix structure likely originating from the human ACE2 region., and about 22% beta sheets, likely to be contributed by the human IgG-Fc adduct. Remaining 32% of the secondary structure of MFcS2 consists of random coils (Figure 3).

**Figure 3:**
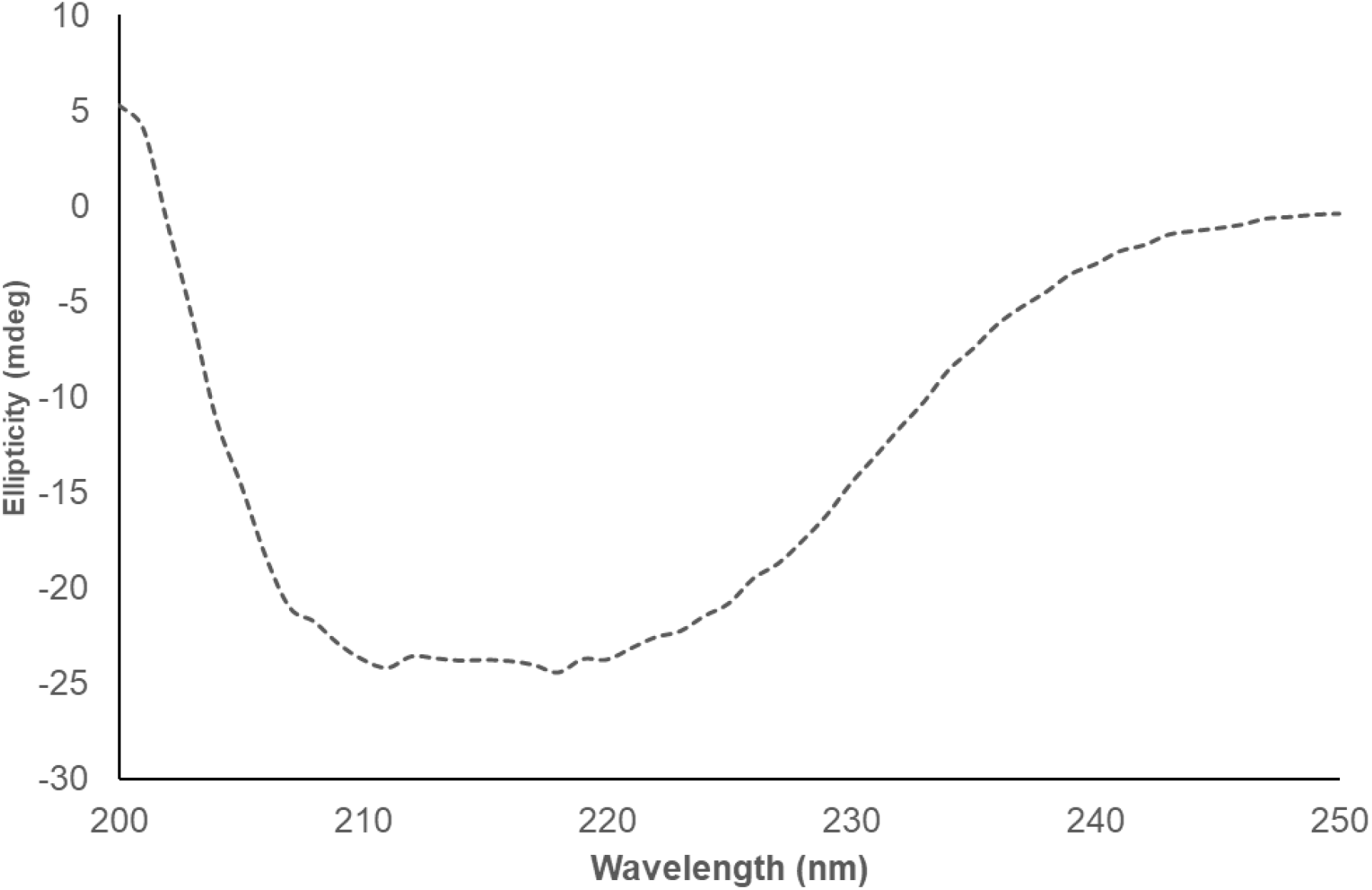
FAR UV CD-Spectra of MFcS2 protein captured in the 200 – 250 nm wavelength range

### 3.3 MFcS2 was enzymatically active

Peptidase activity of MFcS2 was confirmed by its ability to hydrolyze the synthetic fluorogenic substrate Mca=7-methoxycoumarin-4-yl) acetyl;Dnp=2,4-dinitrophenyl [(Mca-APK) (Dnp)]. Incubation of the protein (5 and 2.5 nM) with 100 μM of Mca-APK(Dnp) substrate could successfully cleave the quenched substrate indicated by an increase in fluorescence with time confirming the peptidase activity of the molecule (Figure 4).

**Figure 4:**
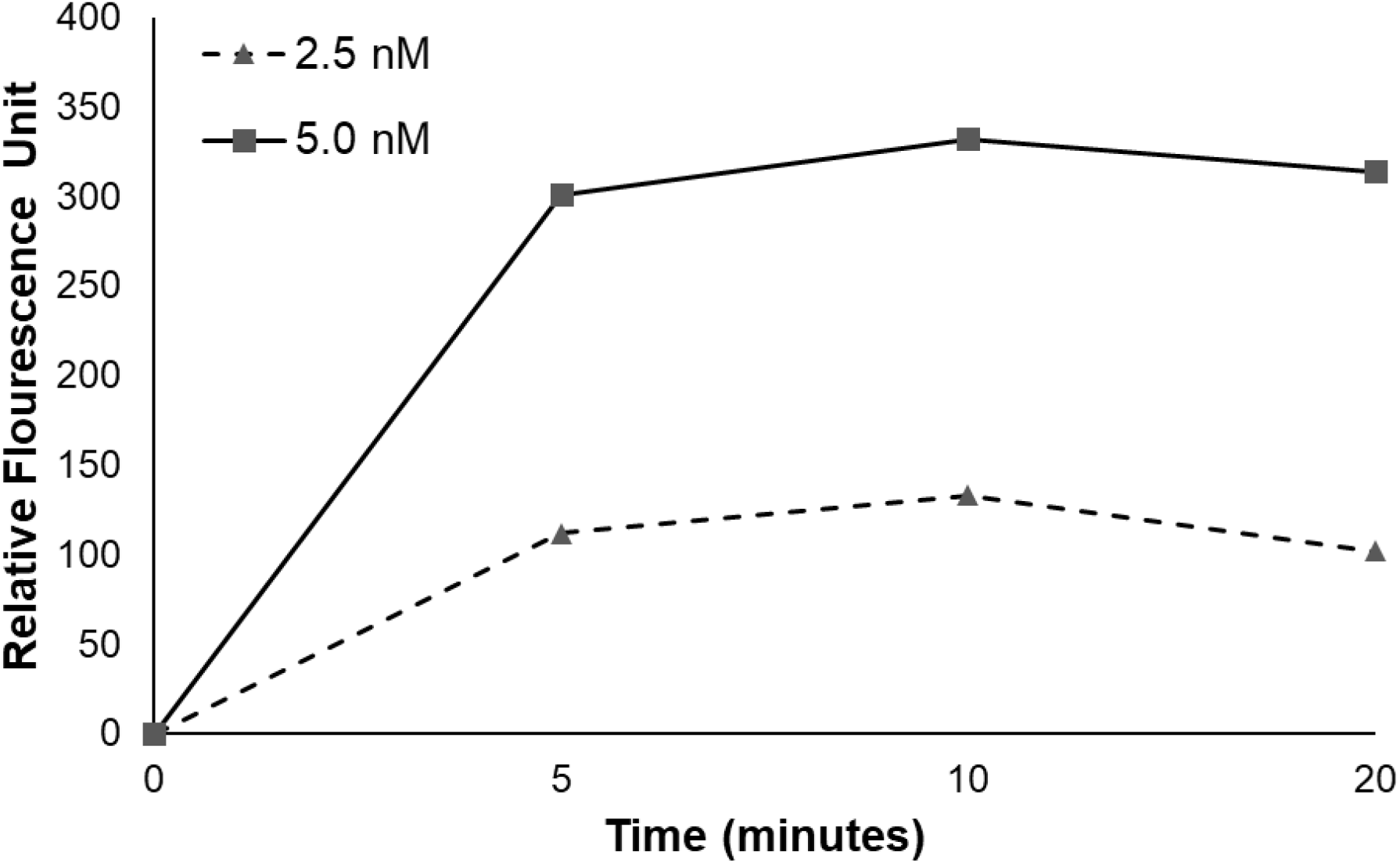
Peptidase activity manifested by two different concentrations of MFcS2 – 2.5nM (triangles with dashed line) and 5.0 nM (squares with solid lines).

### 3.4 MhCT08-E from oral cavity is a potent in vitro model of SARSCoV-2 infection

Transcriptomic analysis revealed 1580 genes differentially expressed (with |log2 Fold Change| ≥2) in infected MhCT08-E cells over mock control, with 311 genes upregulated and 317 genes downregulated (Figure 5a). In agreement with previous studies on viral infection [40], pathways related to defence response to virus and interferon signalling pathways (Figure 5b and c, Table 1) were prominently enriched by the upregulated genes suggesting that the newly described cells from the path of SARS CoV-2 entry, can be used as an in vitro model for studying the effect of the virus.

**Figure 5:**
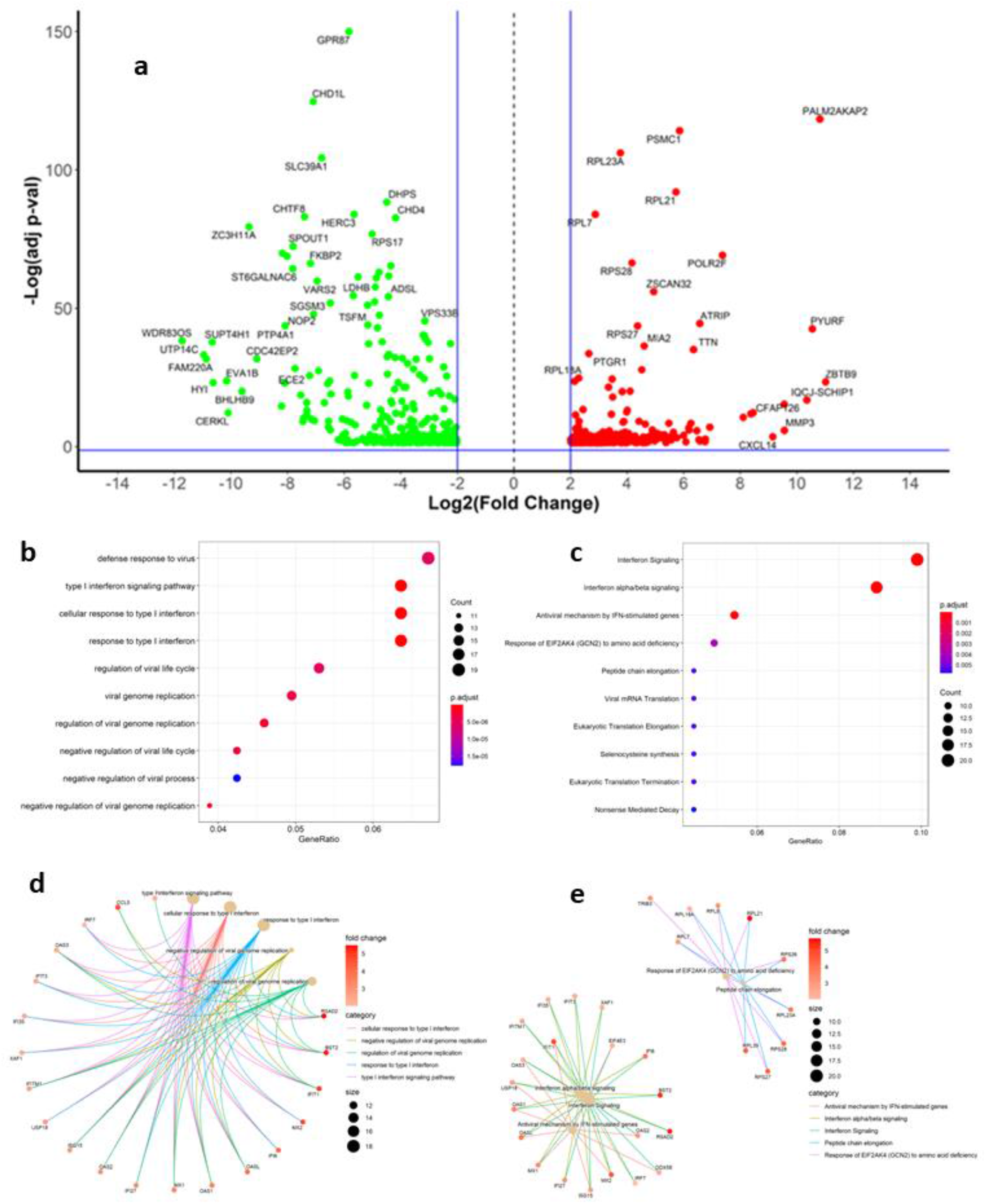
(a) Differentially regulated genes in MhCT08-E cells infected with SARS-CoV2 virus represented as a volcano plot; (b) and (c) GO biological processes and Reactome pathways enriched by the upregulated genes, (d) relationship between upregulated genes and enriched GO biological processes and (e) relationship between upregulated genes and enriched Reactome pathways.

**Table 1:** Upregulated genes and enriched pathways in oral epithelial cells MhCT08 infected with SARS-CoV-2

| Pathways | Gene(s) | Gene names |
| --- | --- | --- |
| **Cellular response to type 1 interferon** | 1. IFIT1<br>2. OAS2<br>3. USP18 | 1. Interferon Induced Protein with Tetratricopeptide Repeats 1<br>2. 2'-5'-Oligoadenylate Synthetase 2<br>3. Ubiquitin Specific Peptidase 18 |
| **Negative regulation of viral genome replication** | 1. OAS1<br>2. OAS3<br>3. OASL<br>4. MX1<br>5. BST2 | 1. 2'-5'-Oligoadenylate Synthetase 1<br>2. 2'-5'-Oligoadenylate Synthetase 3<br>3. 2'-5'-Oligoadenylate Synthetase Like<br>4. MX Dynamin Like GTPase 1<br>5. Bone Marrow Stromal Cell Antigen 2 |
| **Regulation of viral genome replication** | 1. RSAD2<br>2. CCL5<br>3. ISG15<br>4. MX2 | 1. Radical S-Adenosyl Methionine Domain Containing 2<br>2. Radical S-Adenosyl Methionine Domain Containing 2<br>3. ISG15 Ubiquitin Like Modifier<br>4. MX Dynamin Like GTPase 2 |
| **Response to type 1 interferon** | 1. IFITM1<br>2. IFI27<br>3. IRF7 | 1. Interferon Induced Transmembrane Protein 1<br>2. Interferon Alpha Inducible Protein 27<br>3. Interferon Regulatory Factor 7 |
| **Type I interferon signalling pathway** | 1. XAF1<br>2. IFI35<br>3. IFI6 | 1. XIAP Associated Factor 1<br>2. Interferon Induced Protein 35<br>3. Interferon Alpha Inducible Protein 6 |

### 3.5 MhCT08-E grown in high glucose medium showed increased ACE2 and TMPRSS2 expression

MhCT08-E showed overexpression of ACE2 and TMPRSS2 mRNA when grown in higher glucose containing medium (Figure 6) suggesting higher viral entry when compared to cells cultured in lower glucose containing media. This would lead to subsequent cytokine response of cells of oral cavity of individuals with hyperglycemia. This observation is in agreement with the fact that hyperglycemia is a co-morbidity in SARS-CoV2 viral infection [41–43] and the virus gains entry into the host cells through cells of the nose and oral cavity [44] which express ACE2 and TMPRSS2.

**Figure 6:**
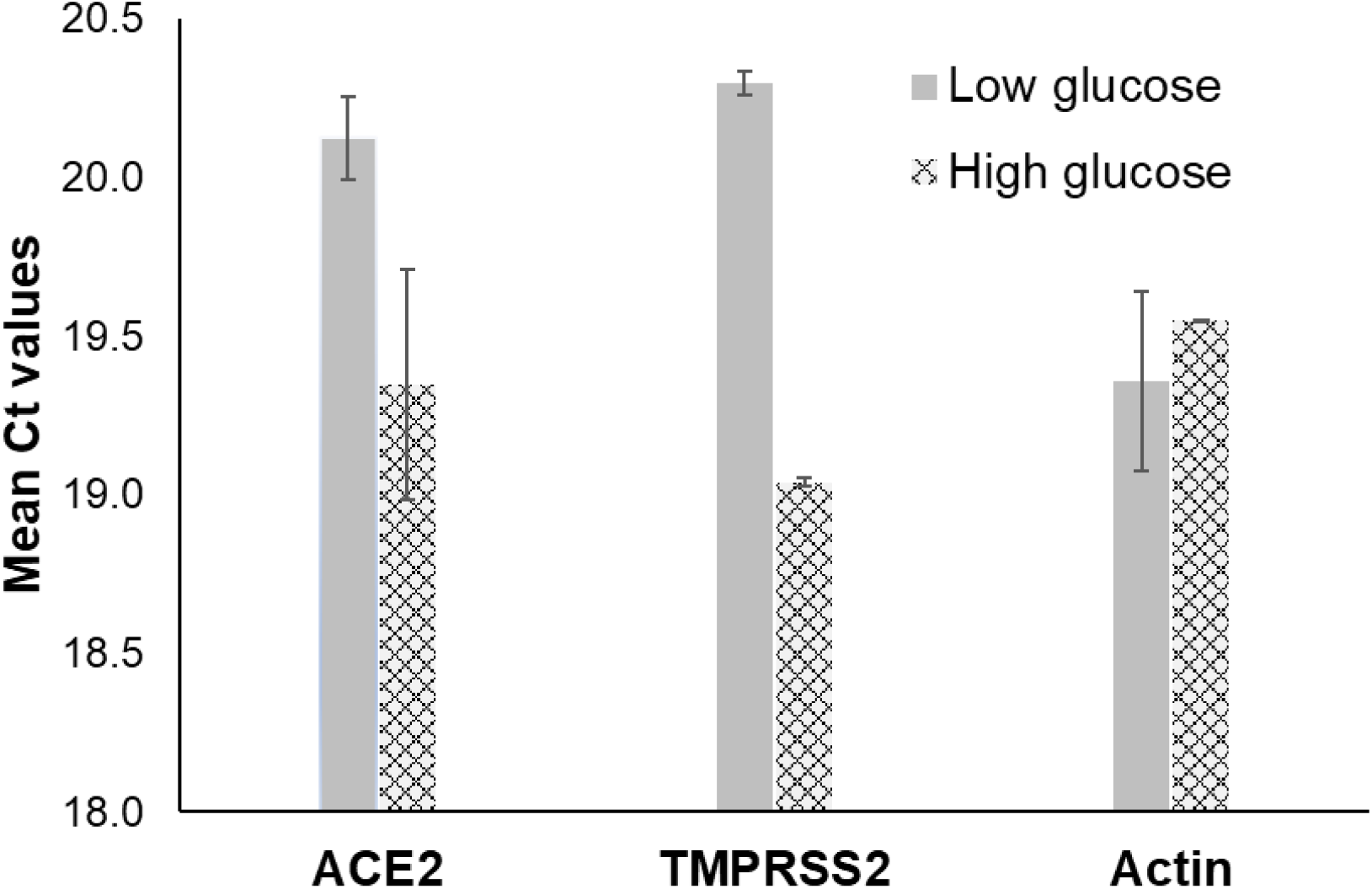
ACE2 and TEMPRSS2 expression in MhCT08-E cells cultured in low (11.1 mM) and high (25mM) glucose medium

### 3.6 MFcS2 showed a dose dependent neutralization of SARS-CoV2 in vitro

LDH enzyme assay results shown in Figure 7 shows nearly 100% cell viability indicting absence of cytotoxicity to Vero-E6 cells treated with MFcS2 at 500, 250, 125, 62.5, 31.25 and 15.625 µg/mL. This suggested that MFcS2 was well tolerated in the cellular system. MFcS2 exhibited a dose dependent percentage reduction in the number of cytopathic plaques in Vero-E6 cells with a maximum reduction of 91.18% at 500 µg/mL and a half maximal inhibitory concentration IC_50_ at 0.608 µM (66.32 µg/mL) (Figure 7). Further, the neutralization potential of MFcS2 was also tested in MhCT08-E and Vero-E6 by qRT-PCR. The protein could neutralize SARS-CoV2 entry in a dose dependent fashion in both cells demonstrated by the reduction in copy numbers of E, N and RdRp genes (Figure 8a and 8b). This suggested a strong binding and neutralization ability of MFcS2, nearly completely abrogating cell penetration at 1.92 uM and 0.48 uM respectively in Vero E-6 and MhCT08-E.

**Figure 7:**
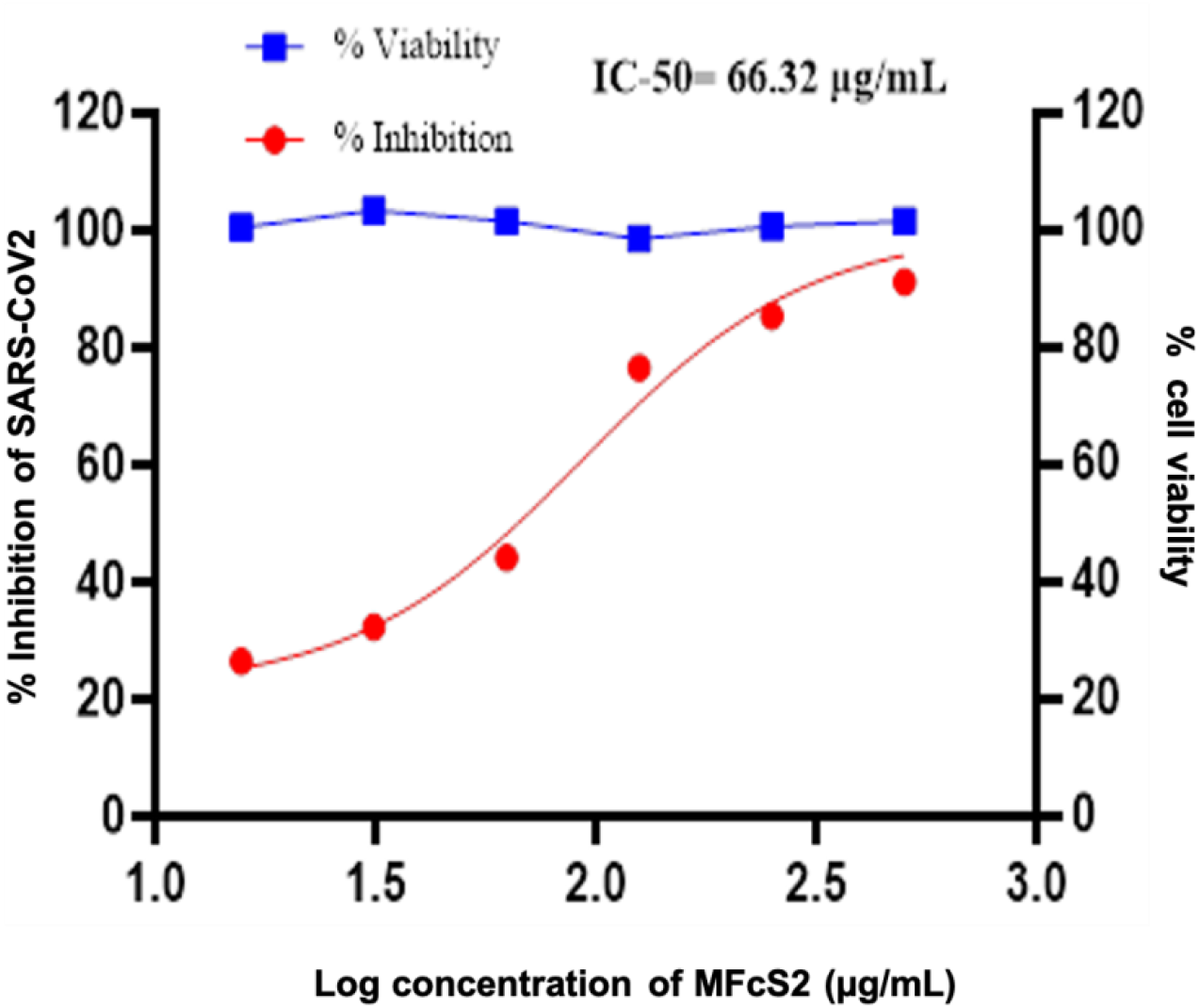
Dose-dependent *in vitro* neutralization of SARS-CoV2 by MFcS2 is observed as %inhibition of cells in PRNT assay (red circles and lines). There is no observed cytotoxicity at these doses of MFcS2 since cell viability is nearly 100% (blue squares and lines).

**Figure 8:**
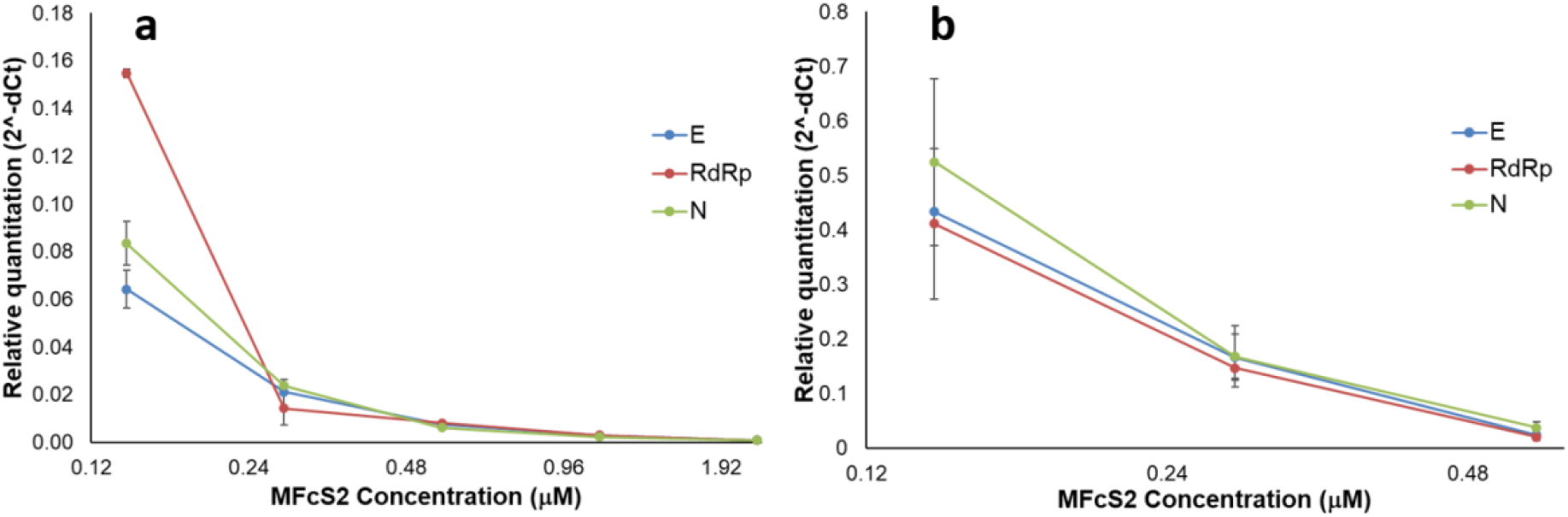
Neutralization of SARS-CoV2 entry to (a)Vero-E6 and (b) MhCT08-E cells by varying concentrations of MFcS2 manifested as relative quantitation of SARS-CoV-2 genes.

### 3.7 MFcS2 exhibited a relatively good half-life in vivo

MFcS2, following IV administration at 5mg/kg distributed rapidly and achieved peak serum concentration of 23.45 µg/mL with a half-life of t_1/2_ of 29.05 hrs (Figure-9a, Table2), whereas MFcS2 delivered intranasally attained peak lung concentration of 8.36 µg/mL with half-life of 11.75 hrs ((Figure9b, Table2). Comparison of computed clearance and AUC values (Table 2) indicates a slower clearance and higher bioavailability of MFcS2 when administered intranasally. This observation indicates the potential therapeutic utility of an intranasal formulation of MFcS2 that would make the drug available for a long period in the lung where maximum pathological effects of SARS-CoV2 has been observed.

**Table 2:** Comparison of pharmacokinetic parameters of a single intravenous dose and a single intranasal dose of MFcS2 in hamsters

| Parameter | Intravenous | Intranasal | Units |
| --- | --- | --- | --- |
| Half life ($t_{1/2}$) | 29.56 | 11.75 | h |
| $T_{max}$ | 0.017 | 1 | h |
| $C_{max}$ | 23453.38 | 8084.06 | ng/mL |
| Clearance | $8.25 \times 10^{-5}$ | $2.37 \times 10^{-5}$ | mL/h/kg |
| Area under the curve | 42498 | 100480 | (ng/mL) × h |

**Figure 9:**
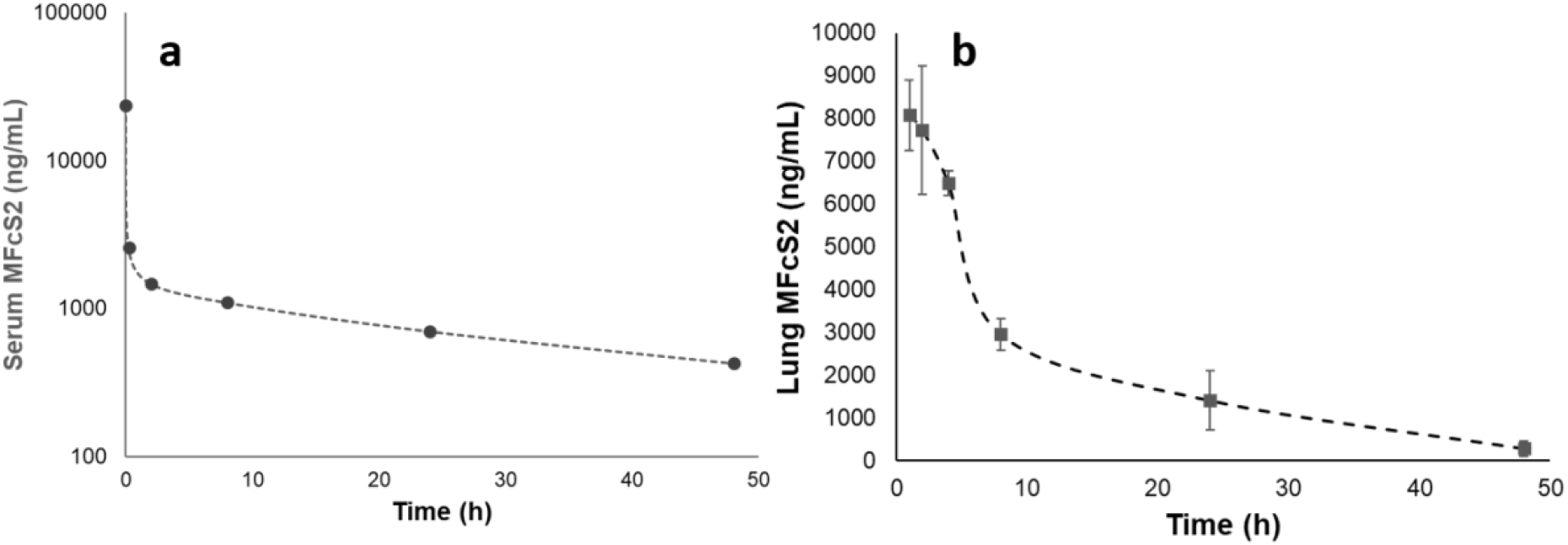
Detected levels of MFcS2 in (a) serum samples after a single intravenous dose and (b) lung homogenate after a single intranasal dose in hamsters. Error bars plotted in both curves but are not visible for serum samples.

### 3.8 MFcS2 did not exhibit any organ toxicity upon in vivo administration

To assess the safety of MFcS2 molecule upon IV and intra-nasal administration, histopathological examination of all vital organs were performed from the hamsters administered with the molecules which revealed normal histoarchitecture in liver, kidney and heart (Supplementary Figure S2-B1;2, D1;2,E1;2) with no gross pathological observations. Spleen from IV administered animal (Figure S2-C1) however showed multifocal areas of neutrophilic infiltration and moderate increase in size of germinal center, whereas appeared normal when administered intranasally (Figure S2-C2). The lung tissue from IV arm showed diffuse mild congestion with multifocal areas of bronchial epithelium showing mild to moderate hyperplasia due to acute lung cellular response to the MFcS2 protein. Interstitial tissue appeared mild to moderately thickened with minimal infiltration of inflammatory cells (Supplementary Figure S2-A1). Lung tissue from intranasally administered animals also showed diffuse mild congestion with multifocal areas of broncho epithelium hyperplasia. In addition, multifocal areas also showed syncytia formation in the bronchial lining epithelium when MFcS2 protein was directly administered intranasally, again possibly due to acute cellular response to the human protein.

### 3.9 MFcS2 exhibited good neutralization potential in vivo when administered therapeutically

The therapeutic potential of the MFcS2 protein was evaluated *in vivo* in hamsters infected with SARS-CoV2 virus and treated with MFcS2 when compared to the untreated group. The viral loads at beginning of day 2 was ∼5×10^6^ PFU that peaked to 10×10^6^ PFU by 12 hrs and then declined to 0.2×10^6^ by day 3 as expected due to self-limiting nature of infection in the Golden Syrian hamster model. MFcS2 treated hamsters group showed relatively reduced SARS-CoV2 PFU numbers in the lungs at each time point (Figure 10a, Treated). Reduction in viral loads/PFU observed in treated group of hamsters ranged from 6-30% at various time points when compared with the untreated group of hamsters, with the highest reduction of 30% observed at 30 hours post infection. The reduction in viral load estimated by PFU was corroborated by increase in cycle threshold (Ct) values of RdRp and N gene in qRT-PCR in the treated group of hamsters (Figure 10b) across all time points. This suggested effectiveness of MFcS2 in neutralizing the virus during the 96 hours treatment period. There was a significant (p<0.01) correlation between SARS-CoV2 detection in lungs of hamsters by PFU vs mean ct values of RdRp and N gene in qRT-PCR (Figure 10c). SARS-CoV2 infected hamsters showed weight loss with the progressing virus infection till day 4, irrespective of whether treated or not. In contrast, the net lung weights increased due to progressive inflammation following infection (Supplementary Table S3).

**Figure 10:**
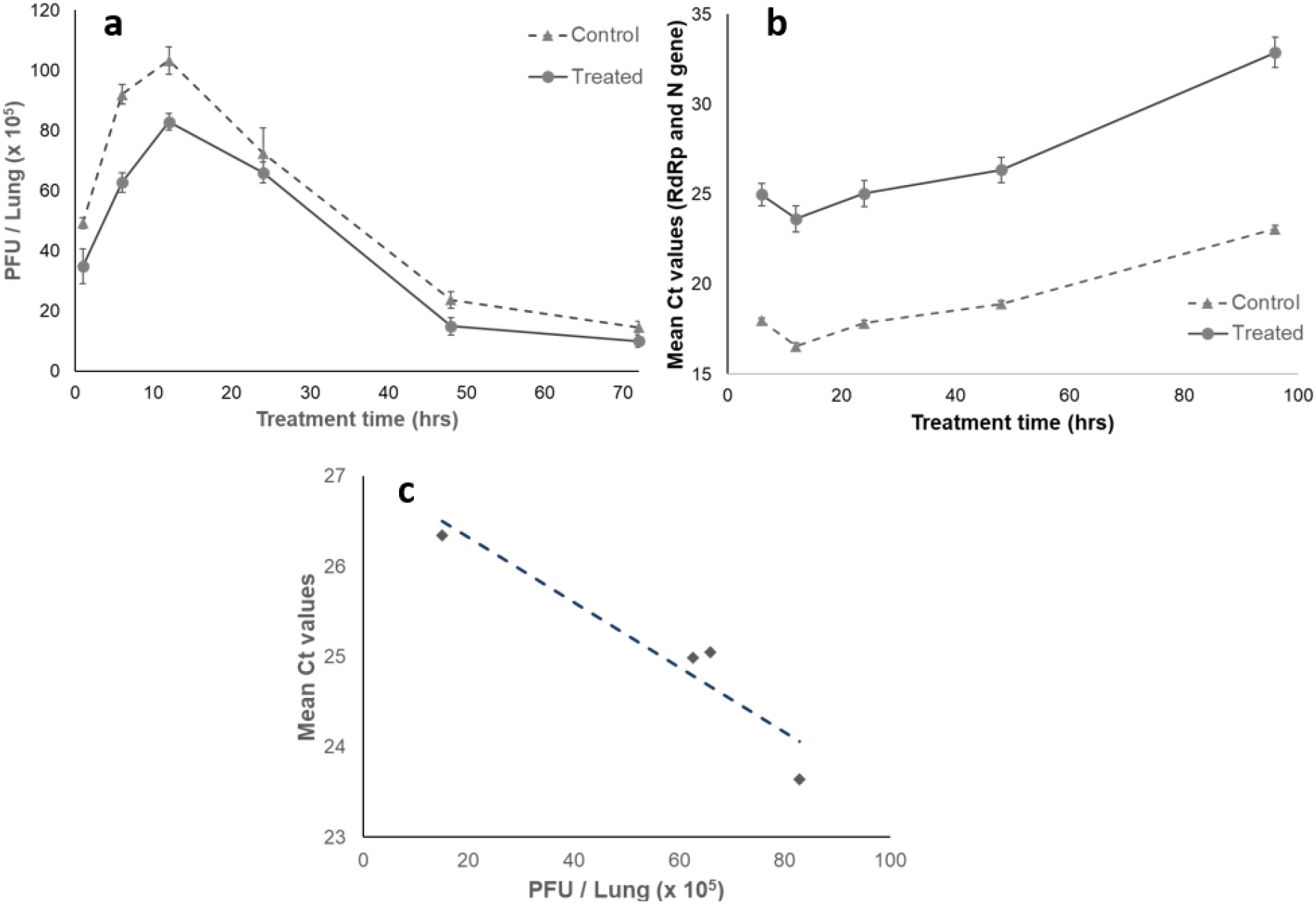
Reduction of SARS-CoV2 PFU in MFcS2 treated hamsters: (a) PFU isolated from lung homogenate from SARS-CoV2 infected but untreated control hamsters at each time point versus PFU from MFcS2 treated hamsters at each time point (b) qRT-PCR Mean Ct values of RdRp and N gene between the control and MFcS2 treated group of hamsters (c) Correlation between PFU count and Ct values in control and MFcS2 treated hamsters.

## 3. Discussion

Multiple therapeutic and preventive strategies have been developed to combat SARS CoV2 infections. To block the binding of SARS-CoV2 spike protein to its cell receptor hACE2 has been considered to be effective by a recombinant soluble ACE2 (rhACE2) which has gone through phase 2 clinical evaluation (NCT04335136) in a small group of people (https://fdaaa.trialstracker.net/trial/NCT04335136). However, the drawback of only using recombinant ACE2 as a therapeutic option is attributed to its limiting serum half-life[45] and studies from multiple groups have shown that an Fc conjugated Ace2 receptor had an improved serum half-life [26, 46]. Fc fusion proteins have prolonged half-life in serum by binding to the neonatal Fc receptor and preventing it from getting degraded. This had led to the use of several therapeutic Fc-fusion proteins in the clinics[47]. Further, structural studies have shown that ACE2 interacts with the receptor binding domain (RBD) of the virus in the form of dimer. A fusion protein with Fc helps stabilize the dimeric structure. Hence, a soluble ACE2-Fc recombinant protein presented in this work is a promising therapeutic candidate for early stage infection of SARS-Cov2[15].

Since the beginning of the pandemic, SARS CoV2 has undergone a series of mutations producing strains with varied morbidity and mortality[48, 49]. One of the mechanisms the more virulent strains have adopted is to mutate the RBD region of spike protein which helps it bind stronger to host ACE2 receptor, thus easing the viral entry into the host system [50–55]. On the other hand several studies have demonstrated the polymorphic nature of ACE2 receptor, altering the susceptibility to the viral infection in individuals [56, 57]. A molecular dynamics simulation study conducted by Wang et al [1] showed that three single-nucleotide variations at K26R, M82I and E329G present in natural variants of ACE2 significantly decreased the binding free energy of ACE2, thus increasing its susceptibility to SARS-CoV2. Further, Fei Ye et al.,[57] showed that N330Y mutation could synergistically be combined with either S19W or T27W to increase the binding affinity. Another study [2] demonstrated that the ACE2-Fc variant with the mutations T27Y, L79T and N330Y in ACE2 was not only expressed and purified with about 80% higher yields but also had higher affinity towards SARS-CoV2 than the wild type ACE2. The same study also demonstrates that engineered Ace2 decoy receptors with T27Y, L79T and N330Y mutations can achieve exceptional breadth against virus sequence variants which could possibly emerge in future[58]. Additionally, a mutational landscape predicted on ACE2 protein in the same study showed that the N90 and N92 residues of ACE2 forms a consensus N-glycosylation motif and all substitutions at these residues, except N92S are favourable for RBD binding. R518G was yet another favourable mutation to increase the binding affinity in the mutation landscape predicted. We thus constructed a modified ACE2 by combining all the favourable mutations (K26R, M82I, E329G, N330Y, R518G,T27Y, L79T, N90Q, N92Q) described above to construct MFcS2 which by design ensures higher binding efficiency with spike protein compared to Ace2-Fc.

Oral cavity which is one of the site of entry of SARS-CoV2 virus also has high amount of ACE2, Furin and TMPRSS2 expression [59]. Thus, in an effort to find an alternate in vitro model of study of SARS CoV2 infection the MhCT08-E cells were shown by gene expression analysis to mimic the molecular profile of viral infection. Further, in line with previous studies [60], MhCT08-E demonstrated an increased level of expression of ACE2 and TMPRSS2 when grown in culture medium with high glucose concentration. Among various pre-existing health conditions [61], diabetes [41–43], hypertension, and chronic kidney disease are the most prevalent comorbidities of COVID-19 known to heighten the risk of fatality [62]. The molecular features of MhCT08-E, thus establish the cell line as a good model of in vitro co-morbidity and viral infection for COVID-19 studies.

Golden Syrian hamster, the model animal used in this study, shares characteristics with SARS-CoV-2 infections in humans by presenting similar lung pathologies [63]. However, the main drawback of this model is that the infection in these animal is self-limiting and they do not usually succumb to the disease [64]. Thus a prolonged efficacy study is not possible with this model. However, this model has been used in multiple studies to demonstrate anti SARS-CoV2 therapies[32, 65]. The efficacy of MFcS2 as a therapeutic candidate was exhibited by up to 30% reduction in functional viral particles isolated from lungs of treated animals in 4 days post infection. The viral load reduction also showed a strong correlation to the reduction in the genome copy number of the virus as demonstrated by qRT-PCR results thus establishing a proof of concept on effectiveness of this molecule as a decoy trap. There are possibilities on further improving efficacy of this molecule by IV dose escalation studies.

Pharmacokinetic analysis following IV route indicated that MFcS2 distributed rapidly from plasma to other body parts. The high volume of distribution suggests MFcS2 is effectively distributed in tissue sites, may be in lungs, the site of infection. The study lacks the information on tissue distribution, following IV dosing, which would be addressed in future. Good PK profile and long half-life of several hours holds promise for achieving *in vivo* efficacy and thus showing its potential as a therapeutic molecule. Slower distribution, a good AUC and absence in plasma of this molecule following intranasal route of administration compared to IV route indicates that MFcS2 remains localized in lungs which is the primary site of infection. May be high levels of expression of neonatal Fc receptors in lung [66], and heart and lung being the primary site of venous circulation localizes the drug to lung. These results thus pave the way to make an intranasal formulation of this molecule which can directly be applied in nasal and oral cavity to block the viral entry. An intranasal spray would require much lesser amount of drug, limiting the side effects, if any, to minimum.

Absence of any histopathological observations of vital organs in the hamsters following MFcS2 administration makes this molecule a promising drug candidate for COVID-19 therapy from safety point of view. However, systematic preclinical toxicology and safety studies are required before progressing further.

In conclusion, MFcS2, has shown promises to be safe and efficacious in relevant *in vitro* and *in vivo* models of SARS CoV2 infection. Being enzymatically active, administration of physiologically feasible amount of the molecule, either through intravenous or intra-nasal route, is expected to bring down the series of events associated with COVID-19 and its comorbidities by maintaining sufficient blood ACE2 levels, allowing proper functioning of RAS pathway [67]. Further systematic evaluations of the intranasal formulation will the next step towards translation of the novel biomolecule to a therapy for early SARS-CoV-2 infection.

## Supplementary Materials

The following supporting information can be downloaded (Figure S1: MFcS2 protein sequence, Table S1: Primers used for constructing the clones, Table S3: Primers used for Ace2 and TMPRSS2 amplification, Figure S2: Histopathology images post IV and IN administration of MFcS2)

## Funding

This research was supported and funded by BIRAC, Grant# BT/COVID0072/02/20

## Acknowledgments

The authors thank Nehanjali Dwivedi and Dr Amritha Suresh MSMF, Bangalore for providing MhCT08-E cell line and Dr Varsha Sridhar of Molecular Solutions Care health for providing the facility to perform the RT-PCR assays.

## Author Contributions

SKD, MD and SPK designed the molecule. SPK performed the experiments and drafted the manuscript. PS and KB helped in the standardisation of transient transfection and assay development. GRM designed and performed invitro studies with VeroE6, MhCT08 and RT-PCR, CNN and BK executed in vivo PK and hamster efficacy studies, RKS and SDN planned and designed invitro, PK and in vivo efficacy studies and preparation of manuscript. BNP and SDS worked on developing the stable cell line overexpressing MFcS2, SS performed qRT-PCR assays, SKD performed the statistical analysis. SD helped in designing the PK assay and critically reviewed the manuscript MD coordinated the study and critically reviewed literature and manuscript.

## Institutional Review Board Statement

All the experimental protocols involving use of animals were reviewed and approved by the IAEC (2082/PO/RC/S/19/CPCSEA) registered with the Committee for the Purpose of Control and Supervision (CPCSEA), Government of India Informed consent statement: Not applicable

## Data Availability Statement

All the data are contained within the article and Supplementary Materials

## Conflicts of Interest

The authors declare no conflict of interest.

## Supplementary Material

**Figure S1:**
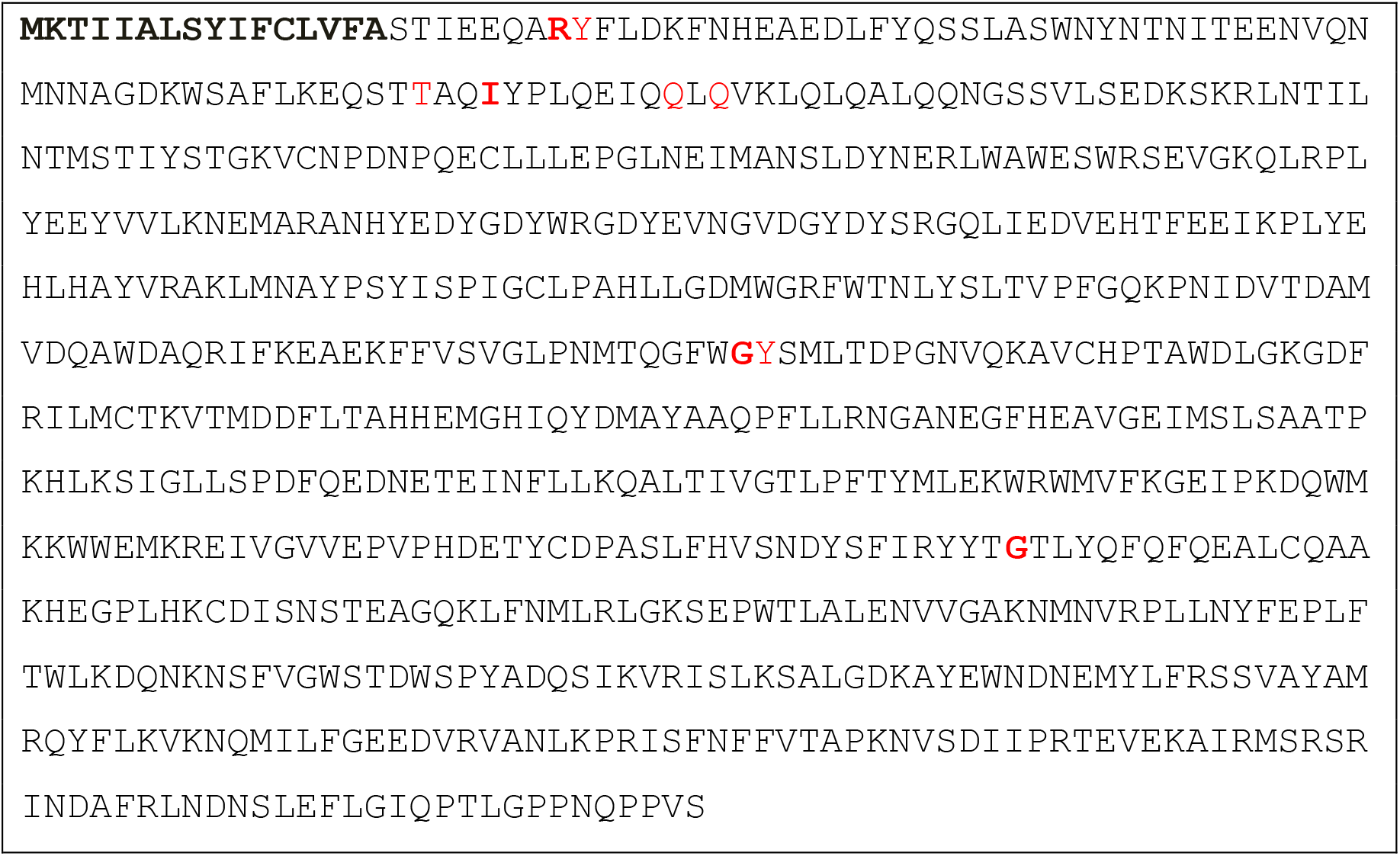
MFcS2 protein sequence

**Table S1:**
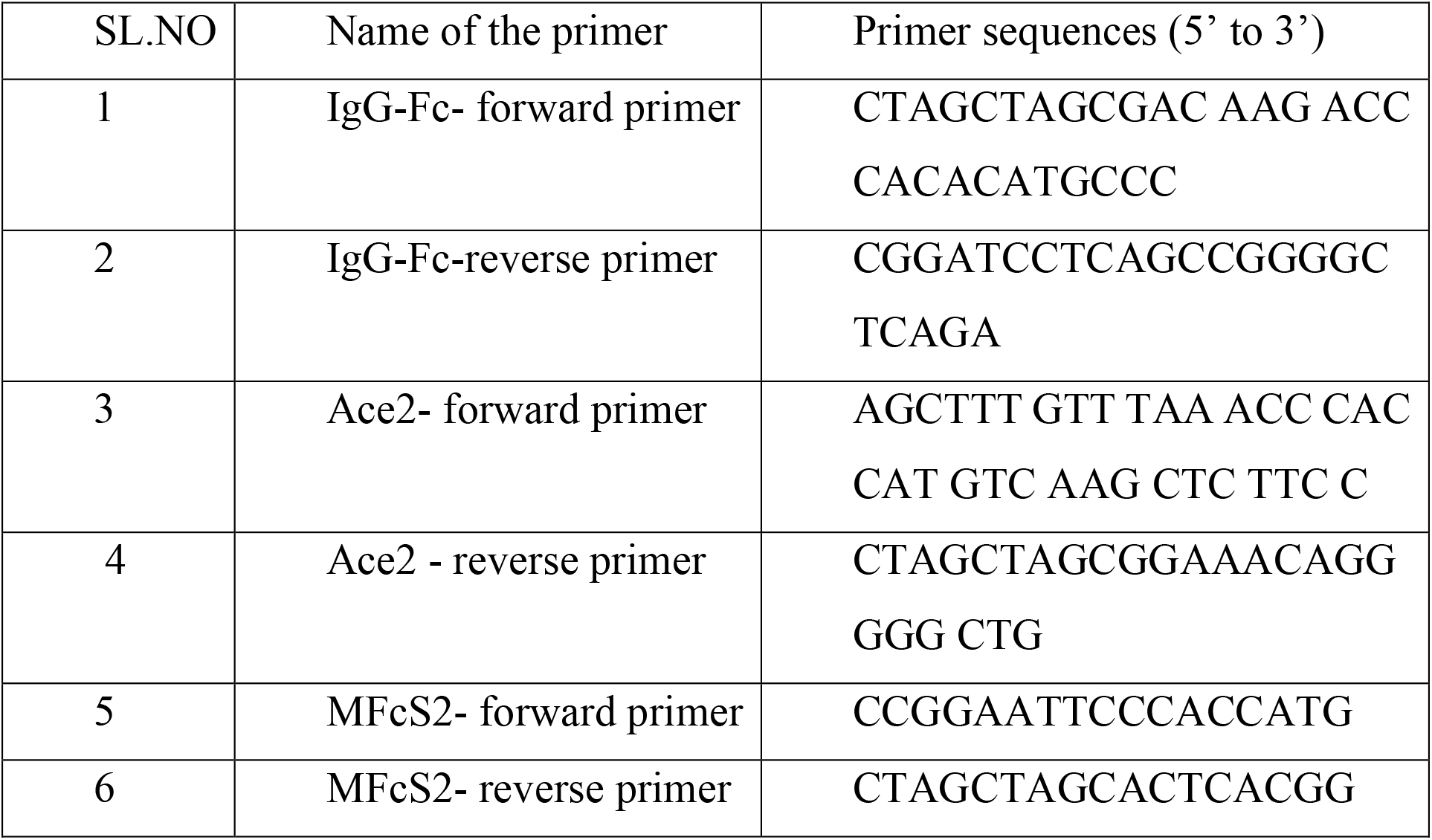
Primers used for constructing the clones

**Table S2:**
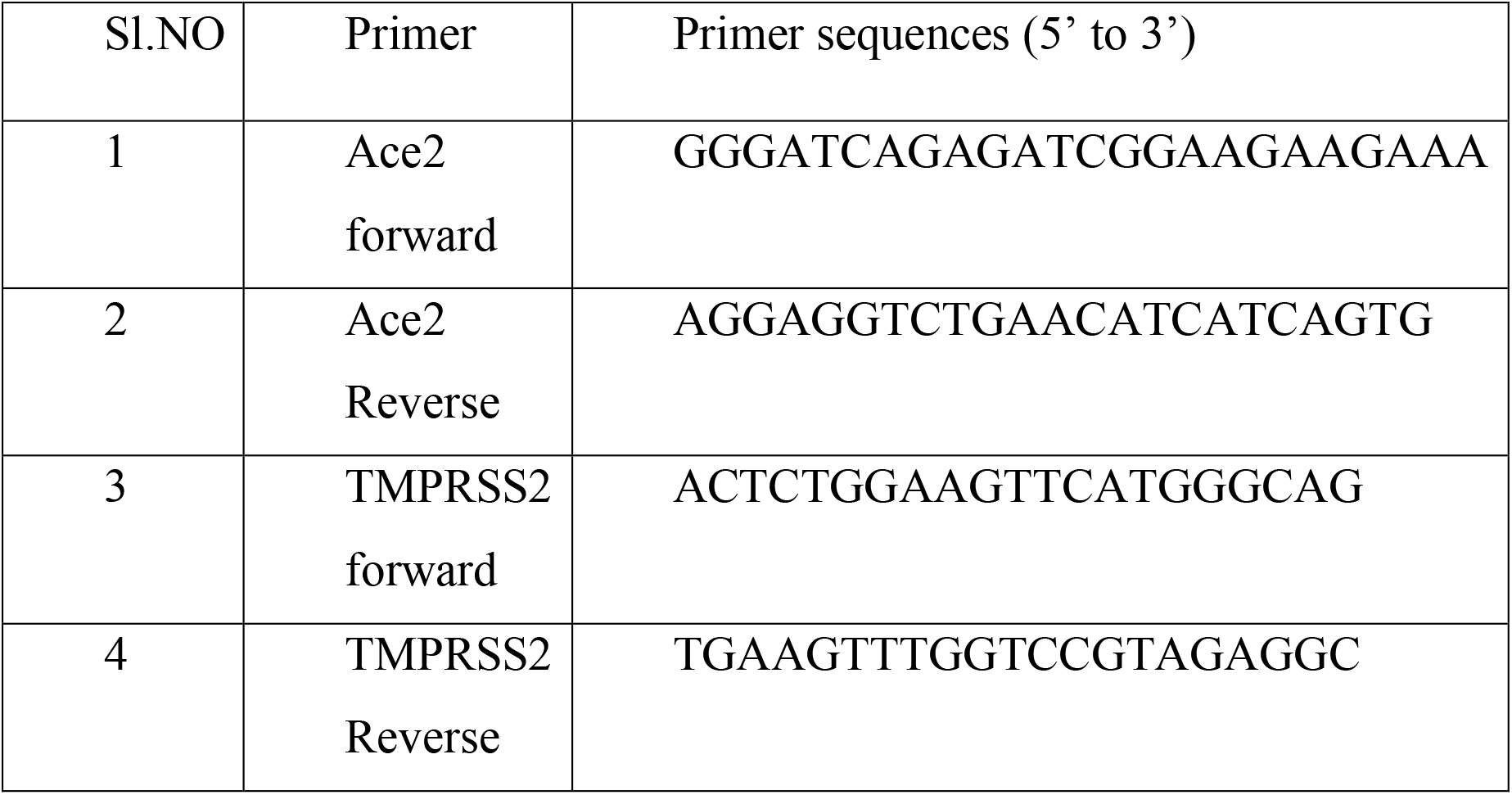
qRT-PCR primers

**Table S3:**
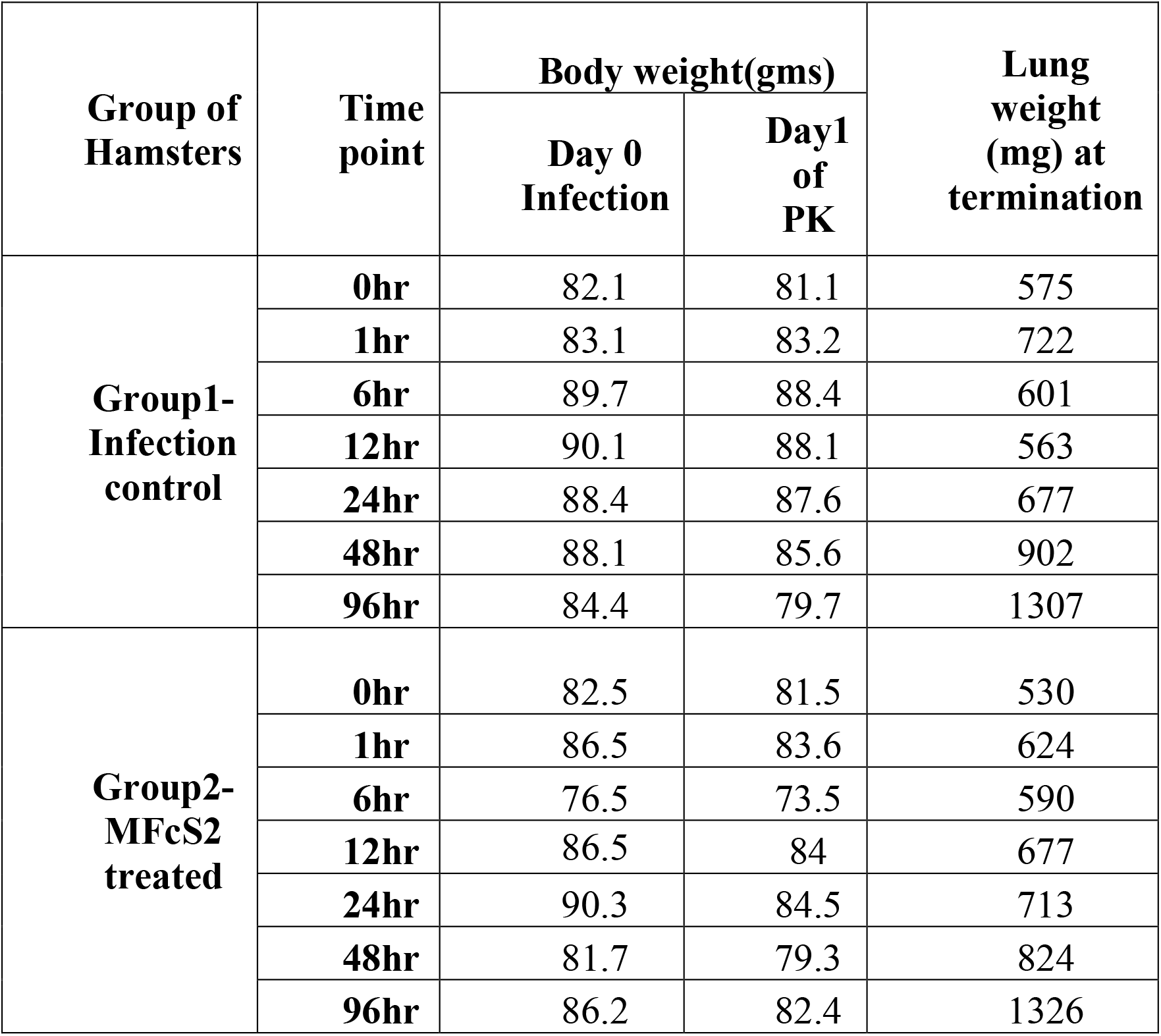
In vivo efficacy study-Hamsters lost weight after SARS-Cov2 infection on day1. Weight of lungs increased after infection due to inflammation. However, increase in weight of lungs in infection and MFcS2 treated lungs were similar.

**Figure 2S:**
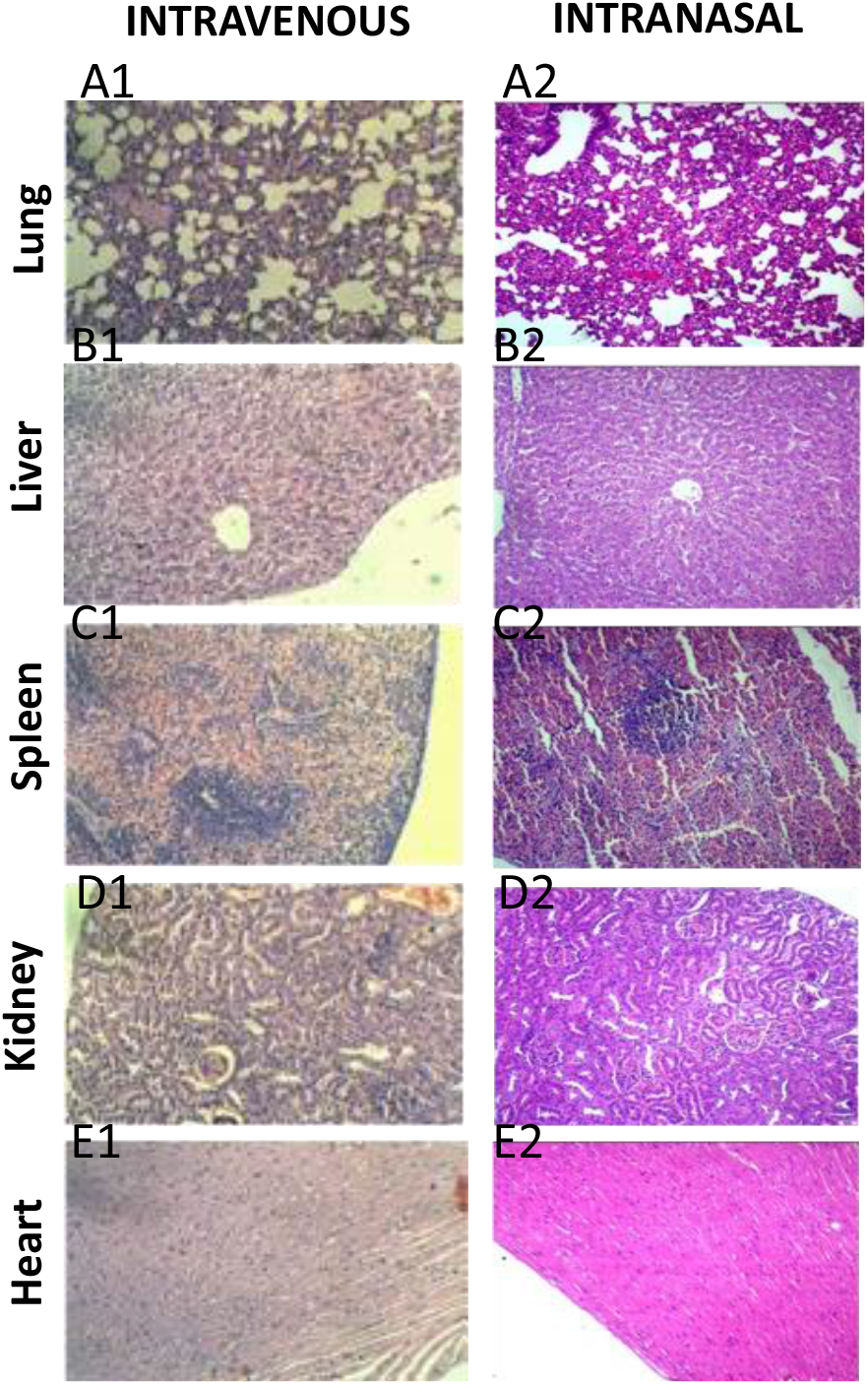
Histopathology findings post IV (A1 to E1) and IN (A2 to E2) administration of MFcS2 protein Intravenous Pathology: (A1) Lungs: The tissue showed diffuse mild to moderate congestion with multifocal areas of bronchial epithelium showing mild to moderate amounts of hyperplasia. The interstitial tissue appeared mild to moderately thickened with minimal infiltration of inflammatory cells, (B1): Liver appeared normal. (C1)Spleen: showed multifocal areas of neutrophilic infiltration and moderate increase in size of germinal center, (D1 and E1) Kidney and heart had no apparent changes. Intranasal Pathology: (A2) Lungs: Lung tissue showed diffuse mild to moderate congestion with multifocal areas of broncho epithelium hyperplasia. The interstitial tissue appeared moderately thickened with minimal infiltration of inflammatory cells. Multifocal areas showed syncytia formation in the bronchial lining epithelium.(B2, C2, D2, E2): Liver, spleen, kidney and heart tissue showed normal histoarchitecture. No other abnormality was observed.

## Notes

### Competing Interest Statement

The authors have declared no competing interest.

